# DNA methylation classification in diffuse glioma shows little spatial heterogeneity after adjusting for tumor purity

**DOI:** 10.1101/2020.03.28.012732

**Authors:** N. Verburg, F.B. Barthel, K.J. Anderson, K.C. Johnson, T. Koopman, M.M. Yaqub, O.S. Hoekstra, A.A. Lammertsma, F. Barkhof, P.J.W. Pouwels, J.C. Reijneveld, A.J.M. Rozemuller, R. Boellaard, M.D. Taylor, S. Das, J.F. Costello, W.P. Vandertop, P. Wesseling, P.C. de Witt Hamer, R.G.W. Verhaak

**Affiliations:** Department of Neurosurgery, Amsterdam UMC, Vrije Universiteit, and Brain Tumor Centre, Cancer Center Amsterdam, De Boelelaan 1117, 1081 HV Amsterdam, The Netherlands; Cambridge Brain Tumor Imaging Laboratory, Division of Neurosurgery, Department of Clinical Neurosciences, University of Cambridge, Addenbrooke’s Hospital, Hill Rd, Cambridge CB2 0QQ, UK; The Jackson Laboratory For Genomic Medicine, 10 Discovery Drive, Farmington, CT 06032, USA; Department of Radiology & Nuclear Medicine, Amsterdam UMC, location VUmc, De Boelelaan 1117, 1081 HV Amsterdam, The Netherlands; UCL institutes of Neurology & Healthcare Engineering, Gower St, Bloomsbury, London WC1E 6BT, United Kingdom; Department of Neurology, Amsterdam UMC, location VUmc, De Boelelaan 1117, 1081 HV Amsterdam, The Netherlands; Department of Neurology, Stichting Epilepsie Instellingen Nederland, Heemstede, The Netherlands; Department of Pathology, Amsterdam UMC, location VUmc, De Boelelaan 1117, 1081 HV Amsterdam, The Netherlands; Department of Neurosurgery, The Hospital for Sick Children, 555 University Ave, Toronto, ON M5G 1X8, Canada; Arthur and Sonia Labatt Brain Tumour Research Centre, Hospital for Sick Kids, Toronto, Ontario Canada; 9Division of Neurosurgery, Li Ka Shing Knowledge Institute, St. Michael’s Hospital, University of Toronto, Toronto, Ontario Canada; Department of Neurological Surgery, UCSF, 505 Parnassus Ave, San Francisco, CA 94143, USA; Princess Máxima Centre for Paediatric Oncology, and Department of Pathology, University Medical Centre Utrecht, Heidelberglaan 100, 3584 CX Utrecht, The Netherlands

**Author notes:** Contributed equally (N. Verburg, F.B. Barthel). Co-senior authors (P.C. de Witt Hamer, R.G.W. Verhaak).

## Abstract

Intratumoral heterogeneity is a hallmark of diffuse gliomas. We used neuronavigation to acquire 133 image-guided and spatially-separated stereotactic biopsy samples from 16 adult patients with a diffuse glioma, which we characterized using DNA methylation arrays. Samples were obtained from regions with and without imaging abnormalities. Methylation profiles were analyzed to devise a three-dimensional reconstruction of genetic and epigenetic heterogeneity. Molecular aberrations indicated that tumor was found outside imaging abnormalities, underlining the infiltrative nature of this tumor and the limitations of current routine imaging modalities. We demonstrate that tumor purity is highly variable between samples and largely explains apparent epigenetic spatial heterogeneity. Indeed, we observed that DNA methylation subtypes are highly conserved in space after adjusting for tumor purity. Genome-wide heterogeneity analysis showed equal or increased heterogeneity among normal tissue when compared to tumor. These findings were validated in a separate cohort of 61 multi-sector tumor and 64 normal samples. Our findings underscore the infiltrative nature of diffuse gliomas and suggest that heterogeneity in DNA methylation is innate to somatic cells and not a characteristic feature of this tumor type.

## Introduction

Diffuse gliomas are the most common malignant brain tumors in adults ^1^. Patients with a diffuse glioma have a poor prognosis and eventually succumb to treatment failure ^2^. The diagnosis, treatment and follow-up of diffuse gliomas rely heavily on imaging ^2^, with magnetic resonance imaging (MRI) as the current standard. Using T1 weighted contrast enhanced (T1c) MRI, diffuse gliomas can be divided into enhancing tumors, predominantly glioblastoma, or non-enhancing tumors, predominantly low grade gliomas (LGG). T1c MRI is used for enhancing and T2/Fluid-attenuated inversion recovery (FLAIR) MRI for non-enhancing gliomas ^3^. However, diffuse glioma infiltration extends beyond the abnormalities detected on standard MRI ^4,5^. Also, the majority of diffuse gliomas recur directly adjacent to the standard MRI-guided surgical cavity ^6^. Heterogeneity of tumor cells is a salient feature of diffuse gliomas and thought to be a driver of treatment failure. Treatment exposure may drive the clonal evolution of heterogeneous tumor cell populations, leading to the selection and survival of resistant subpopulations in some gliomas, whereas refractory disease in others may be driven by other factors ^7^.

Numerous studies have looked at genetic and transcriptomic heterogeneity in diffuse glioma. Recent single-cell transcriptome studies have elucidated transcriptional heterogeneity in regulatory programs that converge on the cell cycle or distinct cellular states ^8,9^ while bulk tissue analysis has demonstrated extensive heterogeneity in somatic drivers such as *EGFR* and *PGFRA ^10,11^* as well as in general somatic alteration burden ^12–14^.

DNA methylation is an epigenetic modification where a methyl-group is added to a cytosine, most commonly measured in the CpG dinucleotide context. These modifications are of interest to the neuro-oncology field as genome-wide patterns in DNA methylation profiles provide a robust method for disease classification and a viable supplement to traditional histopathology ^15,16^. Nevertheless, the extent of intratumoral heterogeneity in DNA methylation remains unclear.

In order to improve our understanding of the (epi-)genetic heterogeneity of diffuse gliomas, we present a comprehensive analysis of DNA methylation of a large number of spatially-separated samples taken from regions with and without imaging abnormalities. We devised a three-dimensional reconstruction of the DNA methylation landscape for each tumor, with particular consideration to the variable ratios of tumor and non-neoplastic cells in each sample. Our analysis demonstrates the infiltrative nature of gliomas beyond visible tumor boundaries with a rather homogeneous DNA methylation landscape across space.

## Results

We analyzed 133 multi-region image-guided stereotactically obtained samples from 16 newly diagnosed and untreated adult patients with a diffuse glioma (Figure 1a, Table 1). Samples were taken from pre-operatively determined sites showing a variable degree of abnormalities. In non-enhancing gliomas, 16 samples were taken outside (FLAIR-) and 41 inside FLAIR abnormalities (FLAIR+). In enhancing gliomas, 12 samples were taken outside both T1c and FLAIR abnormalities (T1c-/FLAIR-), 44 samples outside (T1c-/FLAIR+) and 20 inside T1c abnormalities (T1c+/ FLAIR+) (MRI, Figure 1b). Each sample was profiled for DNA methylation using the Illumina 850k EPIC bead array. Histological slides were stained for hematoxylin-and-eosin (H&E) and MIB-1 and digitized (Figure 1c). Each tumor was reconstructed based on the spatial configuration of the MRI volumes (Figure 2). To reconstruct molecular profiles in three-dimensional space, we calculated the Euclidian distance for each sample to the nearest point on the overall tumor volumes defined by T1c and FLAIR, with negative values indicating points within the tumor volume.

**Figure 1.**
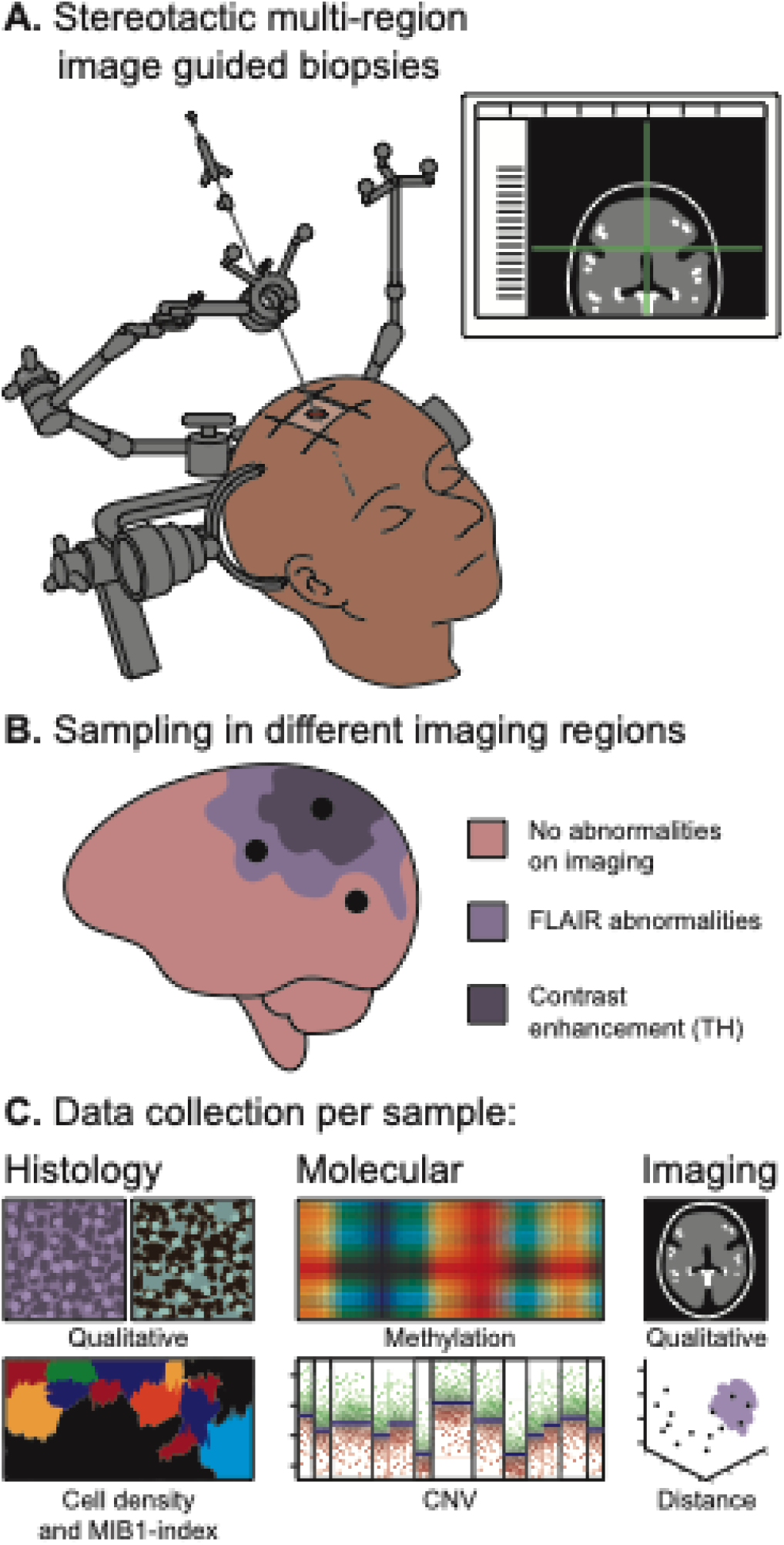
Graphical overview of the methods. **A.** Multiple pre-operatively planned stereotactic biopsies were taken from each patient tumor. **B.** Biopsies were acquired in regions in and outside imaging abnormalities. **C.** Acquired tissue was subject to comprehensive histological and molecular analysis.

**Table 1.** Overview of patients and biopsies in exploration and validation cohort.

| Cohort | Patient | Histology | WHO grade | IDH status | Biopsies |
| --- | --- | --- | --- | --- | --- |
| Exploration | VUmc-01 | Glioblastoma | IV | Mutant | 8 |
| Exploration | VUmc-02 | Glioblastoma | IV | Wildtype | 8 |
| Exploration | VUmc-03 | Astrocytoma | II | Mutant | 7 |
| Exploration | VUmc-04 | Glioblastoma | IV | Mutant | 11 |
| Exploration | VUmc-05 | Oligodendroglioma | II | Mutant | 8 |
| Exploration | VUmc-06 | Astrocytoma | II | Mutant | 8 |
| Exploration | VUmc-07 | Glioblastoma | IV | Wildtype | 9 |
| Exploration | VUmc-08 | Glioblastoma | IV | Wildtype | 10 |
| Exploration | VUmc-09 | Astrocytoma | II | Mutant | 6 |
| Exploration | VUmc-10 | Astrocytoma | II | Mutant | 7 |
| Exploration | VUmc-11 | Glioblastoma | IV | Wildtype | 9 |
| Exploration | VUmc-12 | Astrocytoma | II | Mutant | 8 |
| Exploration | VUmc-13 | Glioblastoma | IV | Wildtype | 9 |
| Exploration | VUmc-14 | Glioblastoma | IV | Wildtype | 7 |
| Exploration | VUmc-15 | Astrocytoma | II | Mutant | 6 |
| Exploration | VUmc-17 | Glioblastoma | IV | Wildtype | 12 |
| Validation | Toronto-01 | Astrocytoma | III | Wildtype | 8 |
| Validation | Toronto-02 | Glioblastoma | IV | Wildtype | 5 |
| Validation | Toronto-03 | Glioblastoma | IV | Wildtype | 7 |
| Validation | Toronto-04 | Glioblastoma | IV | Wildtype | 4 |
| Validation | Toronto-05 | Glioblastoma | IV | Wildtype | 8 |
| Validation | UCSF-01 | Astrocytoma | II | Mutant | 4 |
| Validation | UCSF-04 | Astrocytoma | II | Mutant | 6 |
| Validation | UCSF-17 | Oligodendroglioma | II | Mutant | 3 |
| Validation | UCSF-18 | Oligoastrocytoma | II | Mutant | 4 |
| Validation | UCSF-49 | Oligodendroglioma | III | Mutant | 6 |
| Validation | UCSF-90 | Oligodendroglioma | II | Mutant | 6 |

**Figure 2.**
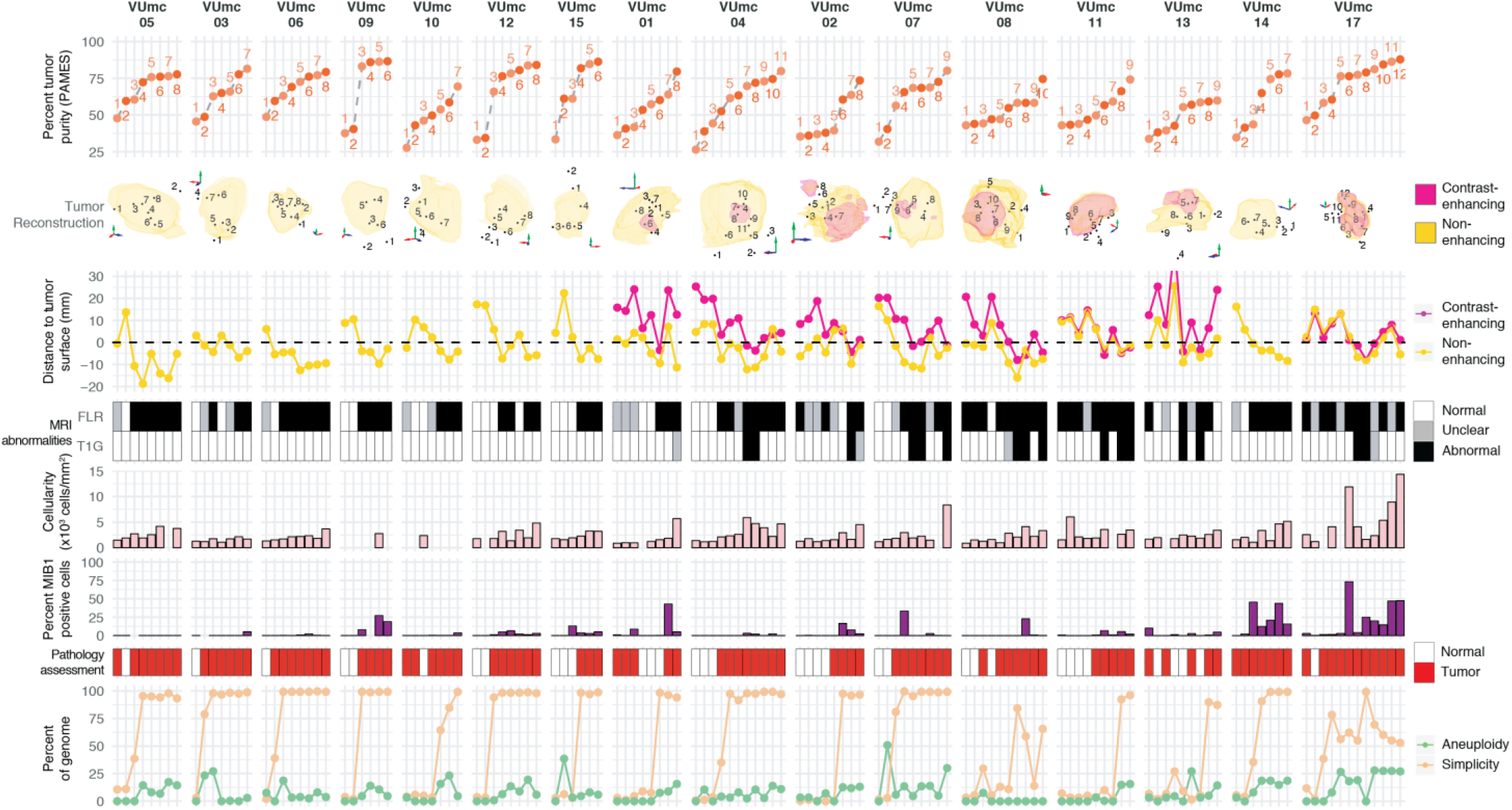
Overview of 133 samples in 16 patients with initial diffuse glioma. Samples are numbered in order of tumor purity for each patient. Imaging-based (distance, FLR/T1G signal) and histology-based (cellularity, MIB1, pathologist classification) parameters are indicated for each sample. PAMES= Purity Assessment from clonal MEthylation Sites.

### Tumor purity is an important determinant of DNA methylation classification and spatial heterogeneity

Since non-neoplastic cells in a sample influence molecular tumor assessment ^17^, we sought to quantify tumor purity, defined as the ratio of tumor to non-neoplastic cells. We evaluated and compared several methods of tumor purity estimation based on histology, MRI, DNA methylation and DNA copy number (Figure 2, Supplementary Figure 1a). DNA methylation-based purity estimates, Purity Assessment from clonal MEthylation Sites (PAMES)^18^ provided the strongest correlations with all other features. Samples from IDH-mutant tumors showed a higher tumor purity than IDH-wildtype tumor samples (Supplementary Figure 1b p=0.03). This was explained by difference in tumor purity between LGG and glioblastoma (Supplementary Figure 1c), with lower tumor purity for the latter due to the known admixture of non-neoplastic cells,^19^ rather than IDH mutational status (Supplementary Figure 1d). The effect of grade on tumor purity is in line with a comprehensive analysis of TCGA samples^20^.

We performed a principal component analysis of the DNA methylation data to elucidate drivers of differences in methylation (258 samples, Figure 3a). Included in the analysis were samples from a second cohort ^16,21,22^ consisting of 61 multi-sector tumor samples from 11 gliomas, and a control cohort of 64 non-neoplastic brain samples. The first principal component (percentage of variance 75%) separated samples based on IDH-status, as indicated by the IDH-mutant and IDH-wildtype samples from the validation cohort (Figure 3a). The second principal component (percentage of variance 4.2%) was driven by tumor purity, as evidenced by the linear increase in tumor purity and the samples from the control cohort (R = −0.69, P <0.01). These findings indicate that tumor purity is the second most important driver accounts for a considerable amount of variation in DNA methylation profiles.

**Figure 3.**
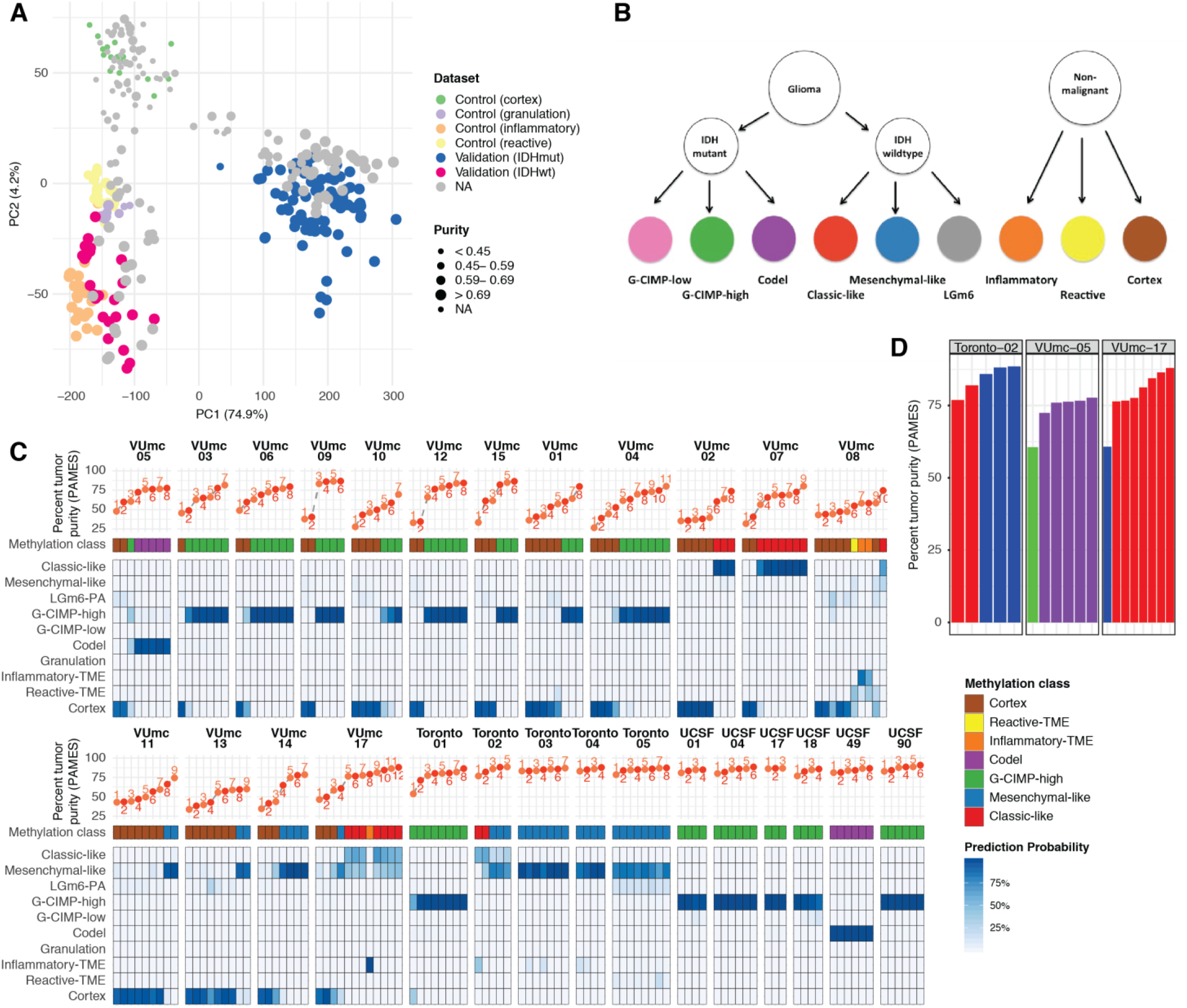
Exploration of spatial distribution of DNA methylation-based subtypes. **A.** Principal component analysis of all (samples=258) samples. **B.** Overview of DNA methylation classification **C.** Overview of DNA methylation subtypes with classification probabilities for exploration and validation cohort. **D.** Bar plot of tumor purity and epigenetic molecular subtype of three patients with spatial heterogeneity.

To establish the relationship between DNA methylation-based subtype and tumor purity, we used L2-regularized logistic regression to assign each sample a tumor subtype, as described in Ceccarrelli et al.,^15^ or non-malignant control subtype (Figure 3b). As expected, samples with a low tumor purity were assigned a control subtype whereas high tumor purity samples were assigned a tumor subtype (Supplementary Figure 2a). The differences in subtype assignment and its relation to tumor purity were clearly captured by the principal component analysis (Supplementary Figure 2b). There were no significant differences between the tumor purity of the different tumor subtypes (Supplementary Figure 2c). Interestingly, the tumor purity of the non-malignant granulation, inflammatory and reactive subtypes was comparable with that of tumor subtypes, which was confirmed by the assessment of all inflammatory and reactive subtype samples as tumor by the neuropathologists. Comparison of the subtypes with another DNA methylation-based classification showed a large conformity for classification families (6.6% discordance, Supplementary Fig 2d) and slightly lower conformity for family subtypes (12.4% discordance, Supplementary Fig 2e).^16^

To explore the heterogeneity of tumor subtypes within a tumor, we analyzed which tumor subtypes were recognized within each patient across the core and validation dataset. The majority of patients (24 of 27) did not show heterogeneity in tumor subtype (Figure 3c). In the three patients with tumor subtype heterogeneity, the one (n=2) or two (n=1) discordant samples were most often the lowest purity tumor sample. Both the Classic-like and Mesenchymal-like subtype were found in two patients, although the high tumour purity samples were Classic-like in one (6 Classic-like and 1 Mesenchymal-like) and Mesenchymal-like (3 Mesenchymal-like and 2 Classic-like) in the other patient. The third patient showed both the Codel (5 higher tumour purity samples) and G-CIMP-low (1 lower tumour purity samples) subtype. These results were confirmed using the classifier from Capper et al. and identify tumor purity as a driving factor of DNA methylation-based tumor subtype heterogeneity (Supplementary Figure 3) ^16^.

### DNA methylation abnormalities extend beyond standard MRI boundaries

To understand the spatial distribution of glioma infiltration, we analyzed the correlation of tumor purity and subtypes with the distance to radiological tumor boundaries and MRI abnormalities. As expected, the distance of samples to the radiological tumor boundaries showed a linear relationship with tumor purity (Supplementary Figure 4a). In non-enhancing tumor, samples classified as cortex were found further away from the radiological tumor boundaries than the other subtypes (Supplementary Figure 4b). Yet, in enhancing tumor this difference was not found, possibly indicating a more diffuse infiltration pattern of enhancing tumors. As anticipated, in enhancing tumors the T1c+/FLAIR+ region showed the highest tumor purity, followed by the T1c-/FLAIR+ and T1c-/FLAIR- (Supplementary Figure 4c). In non-enhancing tumors, tumor purity was higher in the FLAIR+ than the FLAIR-region. Interestingly, samples taken from regions outside standard imaging abnormalities (FLAIR-in non-enhancing and T1c-/FLAIR- and T1c-/FLAIR+ in enhancing tumors) showed a tumor subtype in 36% and 17% of enhancing and non-enhancing gliomas, respectively (Supplementary Figure 4d). Conversely, samples taken from within the standard imaging abnormalities showed a non-malignant subtype in 35% of enhancing tumors, which is most likely due to necrosis in these samples. In non-enhancing tumors 15% of samples within the FLAIR+ region showed a non-malignant subtype. Samples with tumor subtypes were found up to 24mm outside imaging abnormalities. These findings support the diffusely infiltrative nature of these tumors and corroborate the notion that standard MRI does not capture the true extent of diffuse glioma infiltration.

### Spatial heterogeneity of DNA methylation is an innate feature of all somatic cells

To precisely quantify DNA methylation heterogeneity, we performed pairwise comparisons of binarized methylation values between samples. The vast majority of probes were homogeneously methylated (mean 0.93, range 0.83 – 1.0) between samples, suggesting that only a small fraction of probes is responsible for all intratumor heterogeneity. Similar trends have been observed in comparisons of non-neoplastic samples from the same lineage^23^. Unsurprisingly, any two samples from different unrelated tumors showed less probes with identical methylation (mean 0.93 ± 0.02) compared to any two samples from the same tumor (mean 0.96 ± 0.03). However, this difference was subgroup-dependent. For example, any two samples from two unrelated IDH-mutant tumors show more similarity on average than any two samples from two unrelated IDH-wildtype tumors (Figure 4a), likely related to the propensity of (G-CIMP positive) IDH-mutant tumors to uniformly methylate. As expected, a higher degree of heterogeneity can be observed when comparing samples classified as non-malignant to samples classified as tumor, based on DNA methylation classification, within the same patient. Any two IDH-wildtype tumor samples from the same patient show a comparable degree of heterogeneity when compared to two non-malignant samples from the same patient (Kolmogorov-Smirnov *P* = 1.0, pink and green dashed lines). In comparison, any two IDH-mutant tumor samples from the same patient demonstrate less heterogeneity compared to any two non-malignant samples from the same patient (Kolmogorov-Smirnov *P* < 0.0001). These findings suggest that intratumoral DNA methylation heterogeneity is reflecting the clonal nature and shared common ancestor of tumor cells, whereas specimens classified as non-malignant harbor cells from a mixture of lineages.

**Figure 4.**
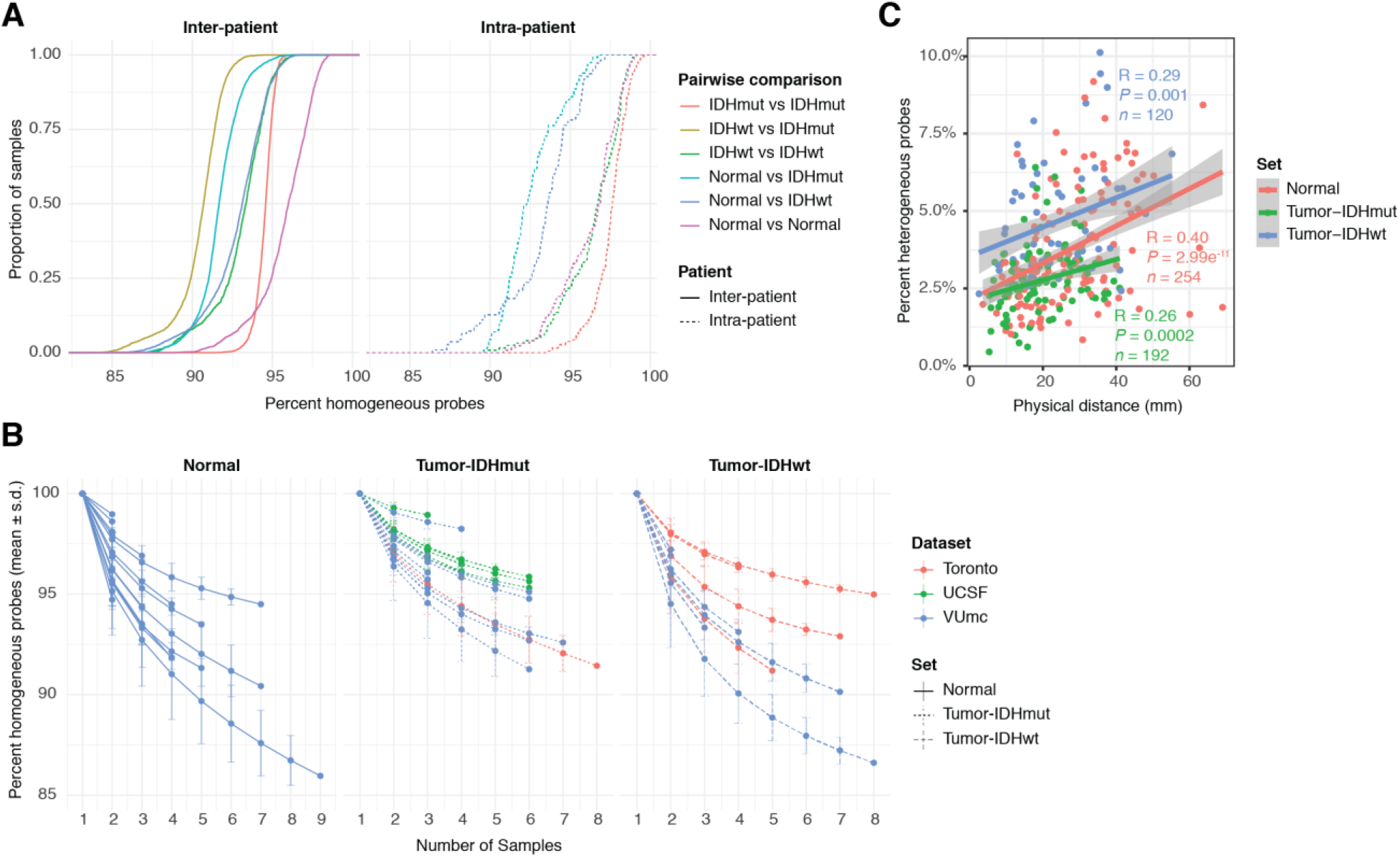
Spatial heterogeneity of genome-wide methylomes. **A.** Empirical cumulative density function (ECDF) curves reflecting similarity (homogeneity) across all pairwise combinations of samples. Comparisons were separated based on whether they involved two samples from the same patient (intra-patient) or between two patients (interpatient) and based on whether the two samples spanned one or multiple sample types. **B.** Line plot showing the cumulative homogeneity associated with additional samples taken from the same tumor. Lines were colored by dataset, tumor and normal samples were separated, and tumor samples were further separated into IDHmut and IDHwt. **C.** Scatter plot of the relation between distance and methylation heterogeneity. Distance is the Cartesian distance in mm between two samples and methylation heterogeneity as described above. Correlation is calculated with Pearson’s R.

To assess the impact of additional samples on tumor heterogeneity we calculated the percentage of identical probes pooling any number of samples per patient, separating samples classified as tumor and normal (Figure 4b). The majority of heterogeneity was captured by the first two samples per patient. Although additional samples further contributed to overall heterogeneity, the change in heterogeneity decreased with each additional sample. Next, we investigated the relation between distance and tumor heterogeneity. These were most strongly correlated in normal samples and less in tumor samples (Figure 4c). This implicates that there is less heterogeneity between two distant tumor samples than between two similar distant normal samples. These patterns were comparable between normal and tumor samples, further supporting the idea that heterogeneity in DNA methylation is largely determined by innate somatic cell (rather than tumor cell) characteristics.

## Discussion

This study represents a comprehensive analysis of spatially-separated samples in diffuse glioma. The combination of histological, radiological and DNA-methylation data enabled us to explore the spatial contexts of tumor purity, epigenetic molecular subtypes and tumor heterogeneity. Our study demonstrates that molecular subtypes are stable and homogeneous after considering tumor purity. Moreover, gliomas are diffusely infiltrative tumors and our data clearly shows that they indeed extend beyond the tumor boundaries found on MRI. Finally, in our study the extent of heterogeneity in tumor cells was predominantly equal to or less than in normal cells.

Information on the spatial heterogeneity of epigenetic molecular subtypes in the literature is limited. A recent study reported intratumor DNA methylation-based subtype heterogeneity in five of twelve glioblastomas in their cohort ^24^. We were unable to confirm this extent of heterogeneity in our study. The differences may be explained by our approach to account for tumor purity prior to determining intratumoral epigenetic subtype classification.

We observed that samples obtained outside imaging abnormalities on FLAIR in non-enhancing and on T1c MRI in enhancing gliomas displayed similar epigenetic molecular subtypes as the core samples. The fraction of tumor cells per specimen varied between the different MRI regions. The presence of tumor tissue outside standard MRI abnormalities is well known ^4,5^. Our results suggest that spatial imaging heterogeneity in glioma is driven by tumor purity, rather than epigenetic heterogeneity. This was confirmed by the strong correlation between tumor purity and the imaging score. Our observations imply that a viable part of the tumor, especially in IDH-wildtype gliomas, is left behind after resection of standard imaging abnormalities.

Tumor heterogeneity has long been viewed as a hallmark of cancer and the idea that various tumorigenic clones compete for resources and evolve in response to treatment pressure is widely accepted. Nevertheless, it is easy to overestimate the role and importance of epigenetic tumor heterogeneity and a more nuanced role for heterogeneity is slowly gaining traction ^25^. Our analysis showed no clear differences in the degree of DNA methylation heterogeneity in tumor tissue compared to non-neoplastic brain tissue, suggesting that a substantial part of the heterogeneity that is detected is innate to somatic cells rather than driven by clonal evolution. In fact, we demonstrated that IDH-mutant diffuse glioma samples show less heterogeneity when compared to surrounding non-neoplastic brain tissue. Nevertheless, it is important to note that the lack of obvious heterogeneity in DNA methylation does not preclude this tumor from demonstrating extensive heterogeneity in gene expression. In conclusion, while diffuse gliomas may show substantial intratumoral heterogeneity at the histological, RNA and protein level, the methylation profile of the tumor cells is rather homogeneous.

## Acknowledgements

N.V. is the recipient of a Dutch Cancer Society Fellowship (OAA/H1/VU 2015–7502) and a Niels Stensen Fellowship. F.P.B. is supported by the JAX Scholar program and the National Cancer Institute (K99 CA226387); K.C.J. is the recipient of an American Cancer Society Fellowship (130984-PF-17-141-01-DMC). FB is supported by the NIHR biomedical research centre at UCLH. We thank Zoe Reifsnyder from the Jackson Laboratory Creative Team for assistance in graphic design. This work was supported by Cancer Center Amsterdam Support Grant 2012-2-05, Cancer Center Support grant P30CA034196; NIH/NCA R01 CA190121, R01 CA237208, NIH/NINDS R21 NS114873, Department of Defense W81XWH1910246 (RGWV).

## Methods

### Sample acquisition and study design

The exploration cohort consisted of 16 patients with an untreated initial diffuse glioma, treated at the Amsterdam UMC, location VU medical center (VUmc), Amsterdam, The Netherlands. Multi-sector sampling was performed, using a stereotactic biopsy procedure preceding the craniotomy, to obtain two samples of each biopsy location for, respectively, FFPE and *Molfix*^©^ (patient 1-8) or snap-frozen (patient 9-16) fixation. The protocol of this study has been published ^26^, was approved by the Medical Ethics Committee of the VUmc and registered in the Dutch National Trial Register (www.trialregister.nl, unique identifier NTR5354). All procedures were carried out in accordance with the Helsinki Declaration ^27^. Written informed consent was obtained from all patients.

The validation cohort comprised 11 patients with 61 FFPE samples from multi-sector sampling of an untreated diffuse glioma treated at the Toronto Western Hospital, Toronto, Canada or USCF Brain Tumor Center, San Francisco, USA. In addition, 64 FFPE samples from 64 patients without a glioma from the German Cancer Network served as controls.

### DNA isolation

DNA isolation was performed by adding proteinase K and incubation at 56°C using QIAamp DNA Mini Kit (Qiagen). DNA was quantified using a Qubit Fluorometer (ThermoFisher). Genomic DNA was bisulfite converted using Qiaamp DNA FFPE tissue Kit (Qiagen)

### DNA methylation profiling by microarray

Data was processed using the minfi packages in R (R Foundation for Statistical Computing, Vienna). Data from the 450k (IlluminaHumanMethylation450k.ilmn12.hg19) and EPIC platforms (IlluminaHumanMethylationEPICanno.ilm10b2.hg19) were processed separately. Detection P-values were calculated for each probe and sample, and samples with an average detection P-value > 0.01 were removed from follow-up analysis. Data was normalized using BMIQ from the wateRmelon package in R. Probes on sex chromosomes and known cross-reactive probes were removed, as were probes mapping to known SNPs and probes with a detection P-value > 0.01. Finally, data from different platforms was merged.

### DNA-methylation based classification and simplicity score

Glioma methylation subtype classification was performed using L2-regularized logistic regression using the R package LiblineaR. Classifiers were trained and evaluated on a set of common probes from TCGA glioma samples with known methylation subtypes. The classes LGm6-GBM and PA-like were merged into a single class LGm6-PA as the separation between these classes was based on phenotype. To improve classification accuracy of samples with low tumor purity, DKFZ controls were added to the classifier as separate classes.

### Methylation purity estimation and simplicity score

DNA methylation measurements of tumor purity included the PAMES algorithm and simplicity score ^17,18^. For the PAMES normal central nervous system samples from the German Cancer Research Center (DKFZ) were used as a control. PAMES operates in three steps. First, AUCs are calculated for each probe discriminating between tumor and normal. Second, a selection of the most informative probes is made. Third, tumor purity is calculated on input samples using these probes

### DNA copy number aberrations inferred from EPIC microarray

Using the R/Conumee package, copy number aberrations were inferred from the 450k and EPIC array data. Merged data from the control samples was used as baseline control for all analyses. Genomic data was used to calculate aneuploidy.

### Immunohistochemistry and qualitative assessment

FFPE samples from the exploration cohort were stained using hematoxylin and eosin (HE) and MIB-1. Two expert neuropathologists independently, blinded for imaging results, assessed the presence or absence of tumor in each sample. Consensus was obtained in case of disagreement. The patient’s histopathological diagnosis was made based on resection material using routine procedures and according to the WHO 2016 criteria.^28^

### Histopathological analysis of whole-slide scans

Using a Hamamatsu Nanozoomer XR, FFPE slides stained with HE and MIB-1 of each sample were digitalized. The 40x magnification images were converted to multiple mosaic images using NDPITools software. Cellularity, defined as number of cells per mm2, was calculated with Cellprofiler. Proliferation index, defined as percentage of Ki-67 positive nuclei of all nuclei, was calculated using local developed software.

### Radiologic evaluation of sample locations

Standard imaging sequences from the patients in the exploration cohort included T1-, T2-, T2/FLAIR and T1c MRI. For each sample location presence of an abnormal signal for each imaging sequence was independently assessed by a neurosurgeon and neurosurgical resident with ample experience in glioma imaging. Consensus was obtained in case of disagreement.

### Sample-to-tumor surface distance

Tumors were segmented on FLAIR and T1c MRI, using Brainlab Software, by a neurosurgical resident with ample experience in glioma imaging. The segmentations and sample coordinates were exported in 3D T1c MRI space and sample to tumor-surface distances were calculated using Matlab.

### Statistical analysis

Median values with interquartile range were used to describe non-normally distribute data. Mann-Whitney-U test was used to compare distributions between subgroups. Correlations were calculated with the Spearman or Pearson’s correlation and compared using Fisher’s z transformation. Comparison of percentages between subgroups was performed using Fisher’s test Normalization and scaling of purity measurement modalities was performed by subtracting the mean and dividing by the standard deviation. To compare absolute purity estimates, the normalized and scaled purity measurements were rescaled using the PAMES mean and SD. P values less than 0.05 was considered statistically significant. R (version 3.5.3) was used for all statistical analyses.

### Heterogeneity analysis

Each probe per patient was classified as methylated (B>=0.3) or unmethylated (B<0.3). A table of all possible pairwise combinations of samples was generated. Each pair of samples was evaluated for heterogeneity by counting the number of identical (homogeneous) probes, the number of differing (heterogeneous) probes and percentages were subsequently calculated. Each pair was annotated according to the metadata for each sample in the comparison.

For each patient and sample type we tabulated all possible combinations of any number samples, iteratively including between 1 and the total number of possible samples. The proportion of heterogeneous and homogeneous probes was calculated when considering each sample in a given set. For each patient/sample type and sample number we then calculated the mean and standard deviation of the proportion heterogeneous across all sets.

## References

1. Ho VK, Reijneveld JC, Enting RH, et al. Changing incidence and improved survival of gliomas. European journal of cancer. 2014; 50(13):2309–2318.

2. Lapointe S, Perry A, Butowski NA. Primary brain tumours in adults. Lancet. 2018; 392(10145):432–446.

3. Ellingson BM, Bendszus M, Boxerman J, et al. Consensus recommendations for a standardized Brain Tumor Imaging Protocol in clinical trials. Neuro-oncology. 2015; 17(9):1188–1198.

4. Pallud J, Varlet P, Devaux B, et al. Diffuse low-grade oligodendrogliomas extend beyond MRI-defined abnormalities. Neurology. 2010; 74(21):1724–1731.

5. Kelly PJ, Daumas-Duport C, Kispert DB, Kall BA, Scheithauer BW, Illig JJ. Imaging-based stereotaxic serial biopsies in untreated intracranial glial neoplasms. Journal of neurosurgery. 1987; 66(6):865–874.

6. Petrecca K, Guiot MC, Panet-Raymond V, Souhami L. Failure pattern following complete resection plus radiotherapy and temozolomide is at the resection margin in patients with glioblastoma. Journal of neuro-oncology. 2013; 111(1):19–23.

7. Barthel FP, Johnson KC, Varn FS, et al. Longitudinal molecular trajectories of diffuse glioma in adults. Nature. 2019; 576(7785):112–120.

8. Neftel C, Laffy J, Filbin MG, et al. An Integrative Model of Cellular States, Plasticity, and Genetics for Glioblastoma. Cell. 2019; 178(4):835–849 e821.

9. Venteicher AS, Tirosh I, Hebert C, et al. Decoupling genetics, lineages, and microenvironment in IDH-mutant gliomas by single-cell RNA-seq. Science (New York, N.Y.). 2017; 355(6332).

10. Snuderl M, Fazlollahi L, Le LP, et al. Mosaic amplification of multiple receptor tyrosine kinase genes in glioblastoma. Cancer cell. 2011; 20(6):810–817.

11. Szerlip NJ, Pedraza A, Chakravarty D, et al. Intratumoral heterogeneity of receptor tyrosine kinases EGFR and PDGFRA amplification in glioblastoma defines subpopulations with distinct growth factor response. Proceedings of the National Academy of Sciences of the United States of America. 2012; 109(8):3041–3046.

12. Sottoriva A, Spiteri I, Piccirillo SG, et al. Intratumor heterogeneity in human glioblastoma reflects cancer evolutionary dynamics. Proceedings of the National Academy of Sciences of the United States of America. 2013; 110(10):4009–4014.

13. Suzuki H, Aoki K, Chiba K, et al. Mutational landscape and clonal architecture in grade II and III gliomas. Nat Genet. 2015; 47(5):458–468.

14. Kim H, Zheng S, Amini SS, et al. Whole-genome and multisector exome sequencing of primary and post-treatment glioblastoma reveals patterns of tumor evolution. Genome research. 2015; 25(3):316–327.

15. Ceccarelli M, Barthel FP, Malta TM, et al. Molecular Profiling Reveals Biologically Discrete Subsets and Pathways of Progression in Diffuse Glioma. Cell. 2016; 164(3):550–563.

16. Capper D, Jones DTW, Sill M, et al. DNA methylation-based classification of central nervous system tumours. Nature. 2018; 555(7697):469–474.

17. Wang Q, Hu B, Hu X, et al. Tumor Evolution of Glioma-Intrinsic Gene Expression Subtypes Associates with Immunological Changes in the Microenvironment. Cancer cell. 2017; 32(1):42–56 e46.

18. Benelli M, Romagnoli D, Demichelis F. Tumor purity quantification by clonal DNA methylation signatures. Bioinformatics. 2018; 34(10):1642–1649.

19. Darmanis S, Sloan SA, Croote D, et al. Single-Cell RNA-Seq Analysis of Infiltrating Neoplastic Cells at the Migrating Front of Human Glioblastoma. Cell reports. 2017; 21(5):1399–1410.

20. Aran D, Sirota M, Butte AJ. Systematic pan-cancer analysis of tumour purity. Nature communications. 2015; 6:8971.

21. Mazor T, Pankov A, Johnson BE, et al. DNA Methylation and Somatic Mutations Converge on the Cell Cycle and Define Similar Evolutionary Histories in Brain Tumors. Cancer cell. 2015; 28(3):307–317.

22. Morrissy AS, Cavalli FMG, Remke M, et al. Spatial heterogeneity in medulloblastoma. Nat Genet. 2017; 49(5):780–788.

23. Ziller MJ, Gu H, Muller F, et al. Charting a dynamic DNA methylation landscape of the human genome. Nature. 2013; 500(7463):477–481.

24. Wenger A, Ferreyra Vega S, Kling T, Bontell TO, Jakola AS, Caren H. Intratumor DNA methylation heterogeneity in glioblastoma: implications for DNA methylation-based classification. Neuro-oncology. 2019; 21(5):616–627.

25. Reiter JG, Baretti M, Gerold JM, et al. An analysis of genetic heterogeneity in untreated cancers. Nature reviews. Cancer. 2019.

26. Verburg N, Pouwels PJ, Boellaard R, et al. Accurate Delineation of Glioma Infiltration by Advanced PET/MR Neuro-Imaging (FRONTIER Study): A Diagnostic Study Protocol. Neurosurgery. 2016; 79(4):535–540.

27. Nathanson V. Revising the Declaration of Helsinki. Bmj. 2013; 346:f2837.

28. Louis DN, Perry A, Reifenberger G, et al. The 2016 World Health Organization Classification of Tumors of the Central Nervous System: a summary. Acta neuropathologica. 2016; 131(6):803–820.

